# Non-canonical mitochondrial STAT3 signaling mediates exercise-induced insulin secretion down-regulation

**DOI:** 10.1101/2021.04.22.440818

**Authors:** Nayara C. Leite, Flávia de Paula, Camila Lubaczeuski, Patricia C. Borck, Jonàs Juan-Mateu, Jane C. Souza, Decio L. Eizirik, Antonio C. Boschero, Jean-Christophe Jonas, Everardo M. Carneiro, Claudio C. Zoppi

**Author notes:** **CORRESPONDING AUTHORS**: Nayara C Leite and Claudio C Zoppi, Obesity and Comorbidities Research Center (OCRC), Department of Structural and Functional Biology, Institute of Biology, University of Campinas. P.O. Box 6109, Campinas, SP. CEP 13083-865, Brazil., E-mail addresses. These authors contributed equally to the study (Nayara C. Leite, Flávia de Paula, Camila Lubaczeuski). Senior co-authorship (Everardo M. Carneiro, Claudio C. Zoppi).

## Abstract

Chronic exercise protects pancreatic beta cells from diabetogenic stress, reducing insulin secretion through unknown mechanisms. We tested the hypothesis that the IL-6/mitochondrial STAT3 (pS-STAT3) axis plays a role in this protective effect. C57BL/6N mice were subjected to endurance training ahead of pancreatic islet isolation and functional analysis. Similar in vitro experiments were performed using insulin-producing INS-1E cells and islets from untrained mice, cultured with serum from trained animals and treated with or without an IL-6 receptor (IL-6R) inhibitor. Then, IL-6R/pS-STAT3 pathway activation and its effects on mitochondrial function and insulin secretion were assessed. Exercise-induced down-regulation of insulin secretion was prevented by inhibition of IL-6R signaling and following STAT3 knockdown. IL-6R activation promoted STAT3 phosphorylation and translocation to the mitochondria, increasing oxygen consumption. Accordingly, lower H2O2 content was reported in islets from trained mice and beta cells exposed to exercise-conditioned serum, while exposure to exogenous H2O2 blocked the down-regulatory effect of training on insulin secretion. Similar findings were observed in islets from obese-trained mice. Together, these findings suggest that the IL-6R/pS-STAT3 axis mediates exercise-induced down-regulation of insulin secretion through modulation of the mitochondrial redox state.

## INTRODUCTION

Chronic insulin hypersecretion caused by congenital defects or related to obesity has harmful outcomes on pancreatic beta cells function and glucose homeostasis (Nessa, Rahman and Hussain, 2016; Bishay and Greenfield, 2016). During the pre-diabetic stages, beta cells increase insulin secretion to maintain normoglycemia in the context of decreasing beta cells mass or insulin resistance. This compensatory hypersecretion is believed to cause beta cell exhaustion, dysfunction and death (Eizirik and Cnop, 2010). Prolonged exercise has beneficial effects for the treatment of metabolic diseases, including improvement of glycemic control in diabetic patients (Jakicic *et al*., 2017). Although this effect has been linked mainly to the increased skeletal muscle insulin sensitivity (Mann *et al*., 2014; Stanford and Goodyear, 2014), there are consistent pieces of evidence showing prolonged exercise modulates pancreatic beta cells function by reducing glucose-stimulated insulin secretion (GSIS) (Calegari *et al*., 2011; Leite Nde *et al*., 2013).

Nevertheless, while the molecular mechanisms promoting an exercise-induced increase in muscle glucose uptake have been deeply investigated, little is known about the mechanisms underlying reduced insulin secretion. By reducing insulin secretion, chronic exercise may protect beta cells from stress and failure, mitigating or preventing high peaks of insulin output. Therefore, unveiling the intracellular mechanisms leading to exercise-induced GSIS down-regulation might support lifestyle interventions while also providing potential targets for the treatment of insulin hypersecretion and diabetes.

Previous work from our group showed that islets from trained rodents produce lower amounts of reactive oxygen species (ROS) (Calegari *et al*., 2012), and that the IL-6/STAT3 axis plays a role in exercise-induced beta cell adaptations (Paula *et al*., 2018). Several studies have shown STAT3 phosphorylation at serine residue 727 (pS-STAT3) enables its translocation to the mitochondria, increasing electron transport chain (ETC) activity and suppressing the production of mitochondrial ROS in different cell types (Meier and Larner, 2014; Yang and Rincon, 2016). Increased ETC activity, leading to lower ROS production and ATP synthesis, has been associated with GSIS down-regulation (Pi *et al*., 2007; Zhang *et al*., 2001). However, the role of mitochondrial STAT3 signaling on ETC activity and its implication for beta cell function has not been investigated. Here we how the IL-6/STAT3 axis is involved in exercise-induced GSIS down-regulation.

## METHODS

All animal procedures were performed according to the ARRIVE guidelines (Kilkenny *et al*., 2010) and to the Principles of Laboratory Animal Care (NIH publication number 85-23, revised 1985), after approval by the local Institutional Committee on Animal Experimentation of the University of Campinas (3057-1) and Université Catholique de Louvain (Project 2013/UCL/MD/9/016).

### Animals

C57BL/6N mice were housed in standard 22°C conditions and kept under specific pathogen-free conditions, in a 12:12h light-dark cycle with free access to food and water in the animal facility of the Health Science Sector of Université Catholique de Louvain or University of Campinas.

At four months of age, male C57BL/6N mice started being subjected to the respective treatments, cited below. To induce obesity, mice received a high-fat diet (HFD) (35% of fat) for 8 weeks. For downstream analysis, one by one, mice were euthanized using a clear CO_2_ chamber. A fill rate of 30% of the chamber volume per minute with carbon dioxide was added to the existing air in the chamber, this rate is appropriate to achieve a balanced gas mixture to fulfill the objective of rapid unconsciousness with minimal distress to the animals accord to American Veterinary Medical Association (AVMA) guidelines.

### Endurance training, IL-6R inhibitor Tocilizumab (TCZ) in vivo treatment, high-fat diet (HFD) protocols and samples collection

Mice were familiarized with a customized treadmill by undergoing 5 mild exercise sessions (at 10 cm/s) for 1 week, before the beginning of the training protocol. The duration of the first session was 10 min and increased 10 min/day. After the familiarization period, mice ran on a treadmill at 70-80% of maximum oxygen consumption (VO_2_ max) with a 20° slope for 1 hour/day, 5 days/week for 8 weeks. Tocilizumab (TCZ, 2 mg/Kg) was injected intraperitoneally, throughout the training protocol. The HFD-treated mice training protocol was initiated along with the HFD diet at 4 weeks of age. After the training protocol period, 48h after the last training session, trained and control untrained mice were euthanized (by exposure to CO_2_, followed by decapitation), and the pancreatic islets and serum obtained from blood centrifugation (4500 rpm at 4°C for 15 min) were collected for additional analysis.

### VO_2_ max test

Immediately before, during, and after the endurance training protocol, VO_2_ max was measured in a sealed metabolic treadmill using a gas analyzer (Oxylet system; Panlab/Harvard Apparatus, Barcelona, Spain), according to a previous protocol (Rezende *et al*., 2006), with the following modifications. The treadmill exercise test was conducted using a 20° slope throughout the test and included a warm-up period of 8 min at 15 cm/s. Subsequently, the treadmill speed was increased by 5 cm/s per minute until exhaustion. Oxygen uptake was recorded at 1-s intervals using specific METABOLISM software (Panlab/Harvard Apparatus) coupled with the gas analyser system. We defined exhaustion as the point when the mice could no longer stay off the shock grid.

### Intraperitoneal glucose test and insulin tolerance test

For the intraperitoneal glucose test (ipGTT), mice were fasted overnight (12 h). An intraperitoneal glucose load (2 g/kg) was administered, and blood glucose measurements recorded at 0, 15, 30, 60, and 120 min via tail snip using a handheld glucometer. For the insulin tolerance test (ITT), mice were fasted for 2 hours and intraperitoneal insulin (Humulin R, Eli Lilly, Indianapolis, USA) load (1 U/kg) was administered. Blood was taken immediately before insulin injection (t = 0 min) and at the times 3, 6, 9, 12, 15, 18, and 21 min via tail snip using a handheld glucometer.

### Islets isolation and culture

Islets were isolated by collagenase digestion of the pancreas (Bordin *et al*., 1995), hand-picked under a stereomicroscope and then cultured at 37°C in the presence of 5% CO_2_ with RPMI 1640 medium (Invitrogen, Carlsbad, CA) containing 10 mmol/l glucose (G10), 5g/L BSA, penicillin and streptomycin.

After isolation, islets recovered *in vitro* for 24h before any experiments, islets were cultured during the recovery period in RPMI 1640 medium supplemented with 10 mmol/l glucose (G10), 5g/L BSA, penicillin and streptomycin. For experiments using mice sera, islets from the control group were incubated in RPMI 1640 medium with 10% serum from control (sCTL) or trained (sTRE) mice for 24h. For experiments using IL-6R antibody, one hour before the treatment with sTRE, islets were treated with TCZ (100ng/mL).

### INS-1E cell culture and treatment

INS-1E cells (passage 71–80; kindly provided by P. Maechler, Centre Medical Universitaire, Geneva, Switzerland) were maintained in RPMI 1640 medium [11 mM glucose, 5% FBS, 1% 4-(2-hydroxyethyl)-1-piperazineethanesulfonic acid (HEPES), 50 µm 2-mercaptoethanol and 1% sodium pyruvate] until 80% confluence. After this period, the cells were pre-incubated with IL-6 (80 ng/ml) for 48h or with the medium containing 10% serum from control or trained mice for 24h.

### Insulin secretion/radioimmunoassay

After the culture treatment period, groups of 5 islets were first incubated for 40 min at 37°C in Krebs-bicarbonate buffer [120 mM NaCl, 4.8 mM KCl, 2.5 mM CaCl_2_, 1.2 mM MgCl_2_, 24 mM NaHCO_3_ and 1g/L of BSA (pH 7.4)], containing 0.5 mM glucose. This solution was replaced with fresh Krebs-bicarbonate buffer, and islets were then incubated for 1 h with 2.8mM or 22.2 mM glucose in the presence or absence of hydrogen peroxide (H_2_O_2_-5µM) with or without TCZ (100 ng/ml).

For determination of insulin secretion, INS-1E cells were incubated for 1h in glucose-free RPMI GlutaMAXI medium and then incubated for 30 min in Krebs-Ringer solution. Cells were then exposed to 2.8mM and 22.2mM for 60 min. Results were normalized by the DNA content measured after cell lysis. At the end of the incubation period, the solution was collected, and insulin content was measured by radioimmunoassay (Scott, Atwater and Rojas, 1981).

### Cellular oxygen consumption

One hour before oxygen consumption (OCR) measurements, the cell medium was replaced by the assay medium (RPMI with no phenol red, 1mM NaPyruvate, 2mM glucose) for 60 min at 37°C (under CO_2_) before loading into a Seahorse XFp Extracellular Flux Analyzer (Billerica, MA, USA). During these 60 min, cartridge ports containing the oxygen probes were loaded with the different compounds injected during the assay and the cartridge was calibrated. Basal respiration was recorded for 30 min, at 5 min intervals, until system stabilization. Glucose was injected at a final concentration of 22 mM, and glucose-stimulated respiration was recorded for 80 min. FCCP was used at final concentrations of 2 µM, Oligomycin 1µM, Antimycin A and Rotenone 1 µM. All respiratory modulators were used at ideal concentrations titrated during preliminary experiments (data not shown), according to manufacturer’s recommendations.

### Hoescht-Propidium Iodide (HO-PI) fluorescence quantification

The percentage of dead and viable cells was assessed using the fluorescence quantification of DNA-binding dyes PI and HO dye 33342 (both Sigma-Aldrich), respectively. INS-1E cells were incubated with HO-PI for 15 min (5 mg/ml) and observed using a confocal laser-scanning microscope (Leica SP5; Leica Microsystems, Buffalo Grove, IL, USA).

### RNA interference

The siRNA for STAT3 was purchased from Sigma Aldrich (NM_012747). The optimal conditions and concentrations of siRNA for beta-cell transfection (30nM) were previously established (Moore *et al*., 2012). Cells were transfected using the Lipofectamine RNAiMAX lipid reagent (Invitrogen) as previously described (Marroqui *et al*., 2014; Nogueira *et al*., 2013). Allstars Negative Control siRNA (Qiagen, Venlo, The Netherlands) was used as the negative control (siCTL). The presently utilized siCTL does not affect beta-cell gene expression or insulin release, as compared with nontransfected cells (Moore *et al*., 2012). After 16 h of transfection, cells were cultured for a 24 h or 48 h recovery period before exposure to serum from control or trained mice.

### Subcellular Fractionation, Western Blotting and immunoprecipitation (IP)

To obtain subcellular fractions, we used the protocol developed by Dimauro et al. (Dimauro *et al*., 2012) with some modifications. Briefly, INS-1E cells were harvested using ice-cold lysis buffer [250 mM sucrose, 1 mM EDTA, 1 mM EGTA, 1.5 MgCl2, 10 mM KCl, 20 mM HEPES (pH 7.5), 1 mM dithiothreitol (DTT), 0.1 mM phenylmethylsulfonyl fluoride (PMSF), 50 mM NaF, 10 mM Na^+^ vanadate, 20 mM Na^+^ pyrophosphate and protease inhibitor cocktail. Cell homogenates were kept on ice for 15 min and then centrifuged at 1000×g for 15 min to sediment nuclear pellets. The resultant supernatant was collected and centrifuged at 11,000×g for 20 min to sediment mitochondrial pellets. The resultant supernatant was collected and precipitated in acetone 100% and then resuspended in STM buffer to obtain the cytosolic fraction. Nuclear pellets were resuspended in NET buffer [20 mM HEPES pH 7.9, 20 % glycerol, 0.5 M NaCl, 1.5 mM MgCl2, 1 % Triton-X-100, 1 mM DTT, 0.1 mM PMSF, 50 mM NaF, 10 mM Na^+^ vanadate, 20 mM Na^+^ pyrophosphate, and protease inhibitor cocktail, whereas mitochondrial pellets were suspended in SOL buffer (50 mM Tris–HCl pH 6.8, 1 mM EDTA, 0.5 % Triton-X-100, protease, and phosphatase inhibitors). Both nuclear and mitochondrial suspensions were kept on ice for 30 min with intermittent shaking and then centrifuged at 9000×g for 30 min to obtain nuclear and mitochondrial fractions, respectively. For Western blotting, 20 µg of the total protein for p-STAT3 Tyr705 (#9131S; Cell Signaling Technology, RRID:AB_331586), α-tubulin (#3873; Cell Signaling Technology, RRID:AB_1904178), p-STAT3 Ser727 (#30647; Abcam, RRID:AB_779085), H3 histone (#9715; cell signaling, RRID:AB_331563), OXPHOS (#ms604; Abcam, RRID:AB_2629281), total STAT3 (#4904; cell signaling, RRID:AB_331269) were resolved using 10% SDS-PAGE and electroblotted onto nitrocellulose membranes. Detection was performed by enhanced chemiluminescence (Pierce). The band intensities were quantified using ImageJ software. For immunoprecipitation, INS1-E cells were treated with IL-6 (80mg/mL) for 48h. The cells were then pelleted and lysed in 500ul of lysis buffer containing 10mM EDTA, 100mM Tris [pH 7.5] and proteinase/phosphatase inhibitors for 40 min on ice. Lysates were centrifuged at 12,000 rpm for 15 min at 4**°**C before immunoprecipitation. The Immunoprecipitated were carried out by incubating 250ug of the total lysate with 5ul of anti-pS-STAT3 on a rotator at 4**°**C overnight. Immunocomplexes were then captured with Protein A agarose (Dynabeads™ Protein A #10001D, Invitrogen). For positive control, the supernatant after incubation with protein A was collected. After three washes with cold lysis buffer, the immunoprecipitated were suspended in Laemlli buffer and separated by Agarose Gel 12%.

### Adenoviruses and Adenoviral infection

Adenoviruses encoding cytosolic and mitochondrial roGFP1 (cyto- and mt-roGFP1) were generated and amplified using the AdEasy XL Adenoviral Vector System (Stratagene, La Jolla, CA) as previously described (Roma *et al*., 2012), and purified on a CsCl gradient. After overnight culture, islets were infected with the adenovirus of interest and treated for 24h with serum from control or trained mice before fluorescence measurements. INS-1E cells were plated, infected 48 h later, and treated with interleukin 6 (IL-6, 80ng/mL) 24 h after infection.

### Dynamic fluorescence measurements

After culture, the coverslip was mounted at the bottom of a 37^°^C temperature-controlled chamber place on the stage of an inverted microscope. Islets or INS-1E cells were perfused at a flow rate of 1 ml/min with a bicarbonate-buffered Krebs solution containing varying glucose concentrations. This solution was continuously gassed with O_2_/CO_2_ (94/6) to maintain a pH of 7.4. Excitation light at appropriate wavelengths was produced by a monochromator: 400/480 nm for -roGFP1, and a 535 nm emission filter was used. The islets were imaged through a ×20 objective and INS-1E cell through a ×40 objective every 30s.

### Statistical Analyses

Data are reported as mean ± SD. Statistical analyses were performed GraphPad Prism (GraphPad Prism version 8, GraphPad Software, La Jolla California USA) using Student’s t-tests, a two-way ANOVA with repeated measures, as appropriate. For two-way ANOVA, if an interaction was found, a post-hoc analysis using the least significant difference-level of statistical significance was set a priori at P < 0.05.

## RESULTS

### Exercise training reduces insulin secretion through IL-6R/STAT3 signaling

Mice were subjected to an eight-week endurance training protocol, displaying by the end of the procedure, typical features of exercise adaptation including increased VO_2_ max and attainment of maximal speed and running distance (Table 1; P < 0.05). Training also reduced fat pads (see supplementary figure 1B-C; P < 0.0001), and increased glucose tolerance and insulin sensitivity (Supplementary Figure 1D-E; P = 0.0237).

**Table 1:**
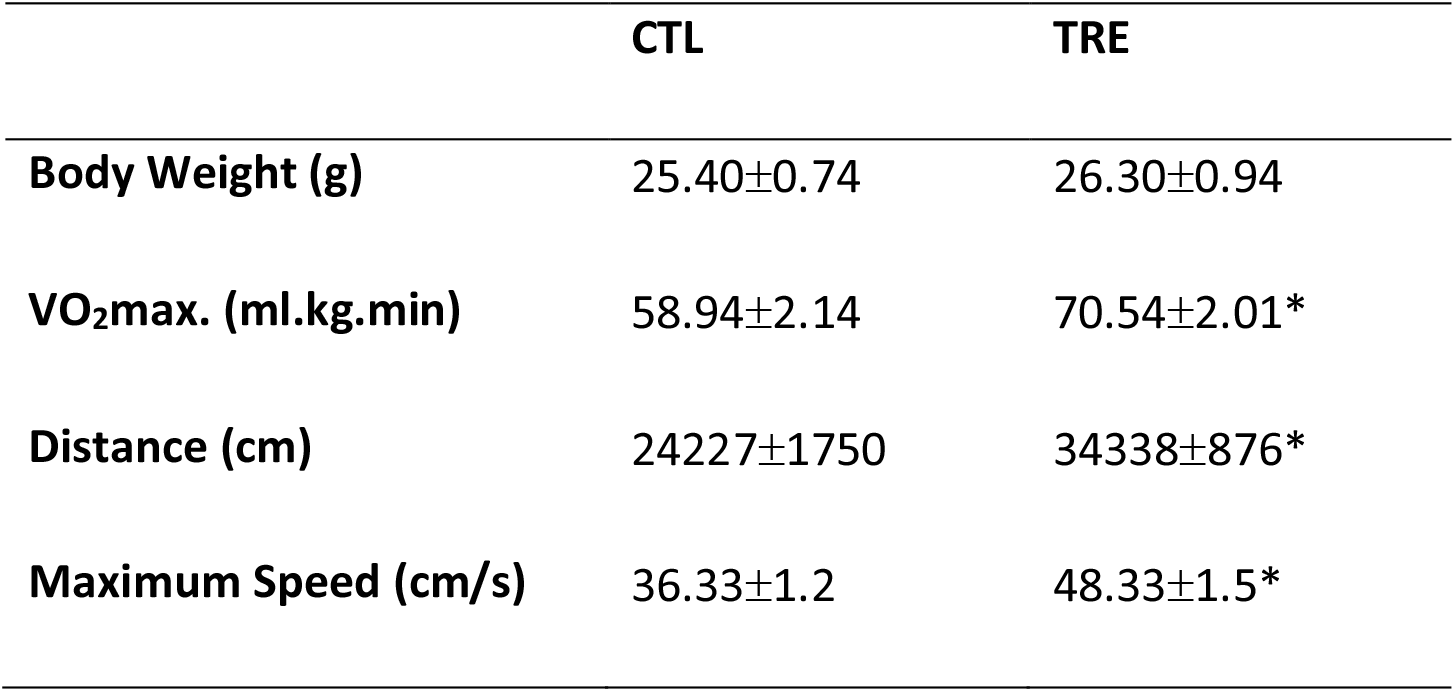
Functional adaptations to the training protocol in C57Bl/6 mice.

|  | **CTL** | **TRE** |
| --- | --- | --- |
| **Body Weight (g)** | 25.40±0.74 | 26.30±0.94 |
| **VO<sub>2</sub>max. (ml.kg.min)** | 58.94±2.14 | 70.54±2.01* |
| **Distance (cm)** | 24227±1750 | 34338±876* |
| **Maximum Speed (cm/s)** | 36.33±1.2 | 48.33±1.5* |

Islets isolated from trained mice displayed reduced GSIS when compared to islets from untrained animals (Figure 1A; P < 0.0001). This effect was blocked when exercised mice were treated with the IL-6 receptor inhibitory antibody TCZ (Figure 1B; P = 0.0204). Similarly, pancreatic islets from untrained mice and INS-1E cells were cultured with sCTL or sTRE, with both groups displayed reduced GSIS when cultured with sTRE (Figure 1C and D; P < 0.001). The exposure to TCZ, one hour before treatment with sTRE, prevented GSIS down-regulation in sTRE cultured islets, suggesting a role for circulating IL-6 in insulin secretion modulation (Figure 1E; P = 0.0181). Using a similar experimental design, we tested whether STAT3 signaling mediates exercise-induced GSIS down-regulation. Knockdown of STAT3 in INS-1E cells prevented the reduction of GSIS induced by the sTRE, confirming the role of STAT3 as a downstream effector of IL-6 signaling (Figure 1F and 1G; P = 0.0139). To strengthen the triggering role of circulating IL-6 on this signaling, we measured IL-6 concentrations in sera. Surprisingly, IL-6 levels were lower in the sera sTRE compared to sCTL (Figure 1H; P = 0.0394). These data provide evidence that exercise training activates IL-6R, regardless of levels of circulating IL-6.

**Figure 1.**
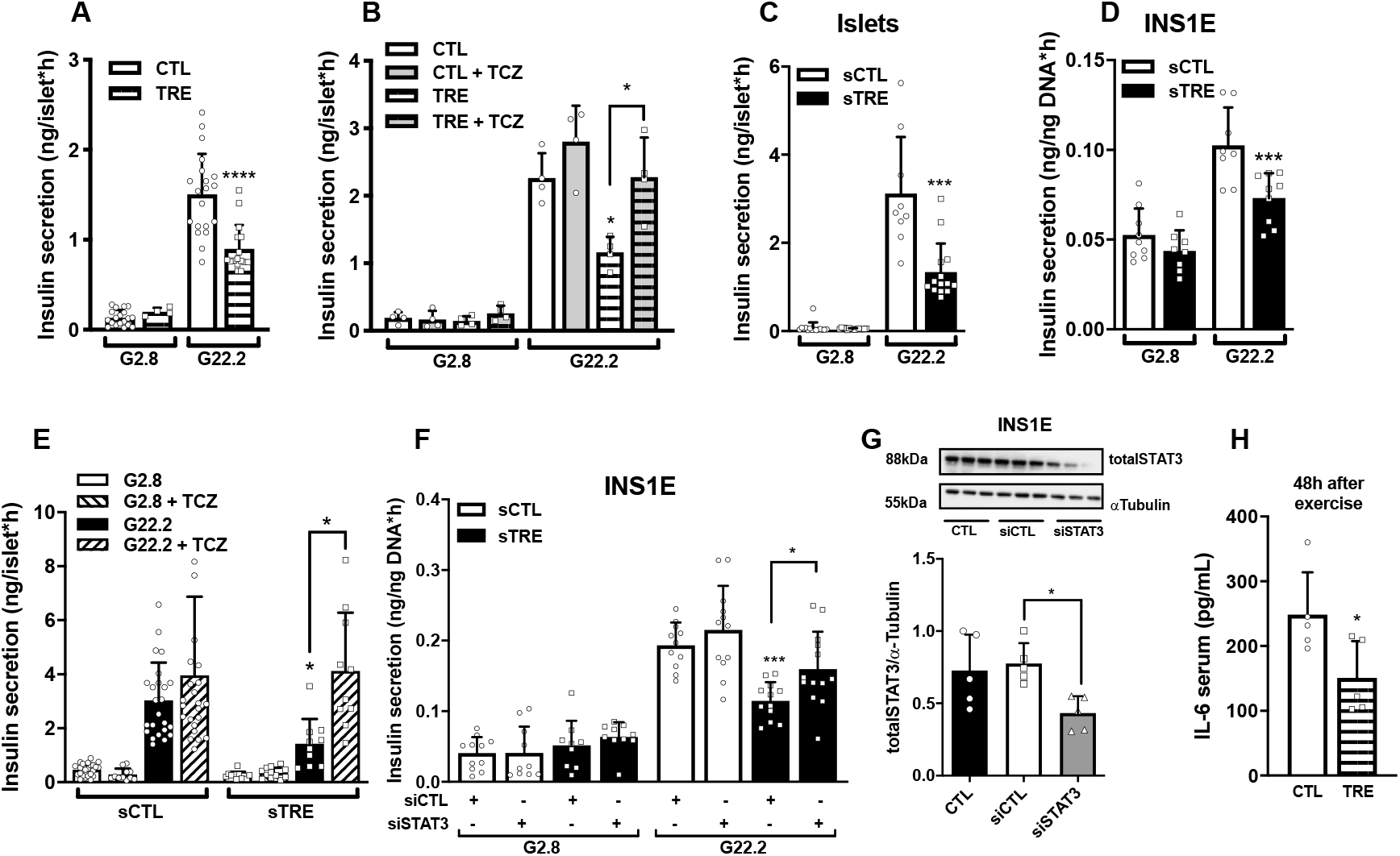
Physical training reduces insulin secretion through IL-6 STAT3 signaling: Glucose-stimulated insulin secretion (GSIS), in response to 2.8mM (G2.8) or 22mM (G22.2) of glucose (A). GSIS in islets from mice treated with TCZ (2mg/kg), injected intraperitoneally daily, during the exercise training period (B). GSIS in islets and INS-1E cells exposed to mice serum during 24h (C, D). GSIS in islets treated with mice serum and TCZ (100ng/mL) during 24h (E). GSIS in INS-1E cells after transfection with a STAT3 siRNA (F). Total STAT3 protein content in INS-1E transfected with siRNA (G). IL-6 sera levels after 48h exercise session (H). Values represent the mean ± SD of three to four independent experiments. *P<0.05/**P<0.01/***P<0.001/**** P<0.0001 vs. Control or as indicated.

### IL-6 signaling triggers STAT3 serine phosphorylation and translocation to the mitochondria

To investigate whether STAT3 translocation to the mitochondria plays a role in the observed phenotype, we sorted mitochondrial, cytosolic and nuclear fractions of INS-1E cells treated with IL-6 and measured the amount of phosphorylated STAT3 in these cell compartments. STAT3 was detected in all cellular fractions. However, phosphorylation of serine 727 (pS-STAT3) was higher in the mitochondria (Figure 2A; P = 0.0033), whereas tyrosine 705 phosphorylation (pT-STAT3) was more prevalent in the nucleus (Figure 2B; P = 0.1183). To confirm fraction purity, specific markers of protein content for each cellular fraction were quantified (Supplementary Figure 2A). Higher content of pS-STAT3 was also observed in islets from trained mice (Figure 2C, P = 0.0153), supporting the role of exercise as a stimulus for mitochondrial STAT3 translocation. To test whether STAT3 phosphorylation is mediated by IL-6R signaling, we treated mice with TCZ along with training, and pre-incubated INS-1E cells with TCZ prior to the exposure to sera of trained mice (sTRE). Inhibition of IL-6R signaling both *in vivo* and *in vitro* prevented the increase of pS-STAT3 expression (Figure 2D and E; P = 0.037), supporting the role of IL-6R on STAT3 serine phosphorylation and its subsequent translocation to the mitochondria. In line with this, immunoprecipitation of pS-STAT3 and western blotting showed that binding of pS-STAT3 to ATP synthase (complex V) is increased in INS-1E after IL-6 exposure (Figure 2F and G; P = 0.0165). The treatments used in this study, including sera, IL-6, and downregulation of STAT3 did not affect beta cell viability (Supplementary Figure 2 B-D, P < 0.0001).

**Figure 2.**
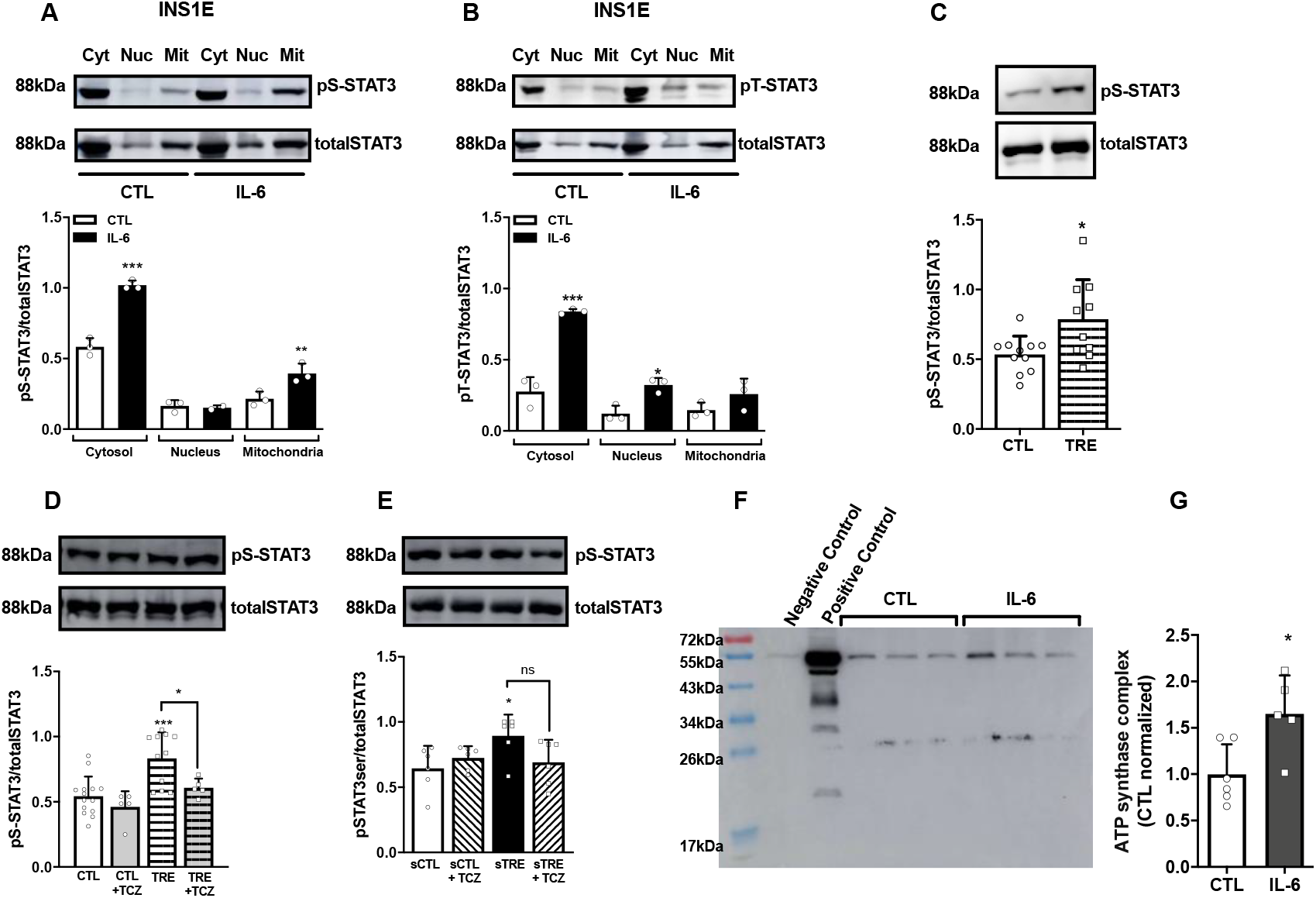
IL-6 modulates beta cell function through pS-STAT3: Protein content in IL-6-treated INS-1E cells of total STAT3, Ser727 STAT3 (pS-STAT3) (A) and Tyr705 STAT3 (pT-STAT3) (B) in cytosolic (Cyt), nuclear (Nuc) and mitochondrial (Mit) fractions. Protein content is shown for total STAT3 and pS-STAT3 in islets from control (CTL) and trained (TRE) mice (C). Protein content is shown for total STAT3 and pS-STAT3 in islets from control (CTL) and trained (TRE) mice treated with TCZ (1mg/kg) during the exercise training period (D) and in islets from untrained mice treated for 24h with serum from control (sCTL) and trained (sTRE) mice treated with TCZ (100ng/mL) (E). INS-1E cells were treated with IL-6 for 48h, followed by immunoprecipitation of pS-STAT3. pS-STAT3 precipitates were subjected to immunoblotting for OXPHOS (F). Quantification of ATP synthase complex bound to pS-STAT3 (G). Values represent the mean ± SD of three to eleven experiments. *P<0.05/**P<0.01/***P<0.001/****P<0.0001 vs. Control or as indicated.

### Chronic exercise increases ETC activity and reduces ROS in beta cells

We evaluated whether IL-6R/STAT3 axis signaling affects ETC activity in beta cells by measuring oxygen consumption. After exposure to IL-6, INS-1E cells displayed higher levels of maximal respiration (response to FCCP following oligomycin). By knocking-down STAT3, the signaling of the IL-6R/STAT3 axis on this functional outcome was strengthened. The lack of STAT3 reduced oxygen consumption rates (Figure 3A; P = 0.2916). Next, we analyzed the impact of training on ROS production and ATP flow. Using the amplex Ultra Red probe, which provides an overview of the H_2_O_2_ content, we found islets from trained mice displayed lower amounts of ROS production (Figure 3B; P = 0.0067). To confirm the modulation of the beta cell redox state by prolonged training and IL-6R/STAT3 axis activation, we transduced mouse islets and INS-1E cells with both cytosolic and mitochondrial redox-sensing green fluorescent proteins (roGFP1). Although we were unable to detect significant changes in the cytosolic thiol oxidation of islets cultured with sTRE (Figure 3C-D), IL-6 exposure significantly reduced cytosolic thiol oxidation in INS-1E cells (Figure 3E-F; P = 0.001). The roGFP1 targeted to mitochondria (mt-roGFP1) showed a slight but significant decrease in thiol oxidation in islets cultured with sTRE and INS-1E cells cultured with IL-6 (Figure 3G-J; P = 0.0323), evidencing lower mitochondrial ROS production. Surprisingly, ATP flow was not altered (Figure 3K-L, P = 0.8676). To confirm the role of enhanced ETC activity with consequent lower ROS content on GSIS down-regulation, we measured GSIS in the presence of exogenous H_2_O_2_. Islets from trained mice, as well as islets from untrained mice cultured with sTRE, lost the down-regulatory effect of training on GSIS in the presence of exogenous H_2_O_2_ (Figure 3M, N; P < 0.0001).

**Figure 3.**
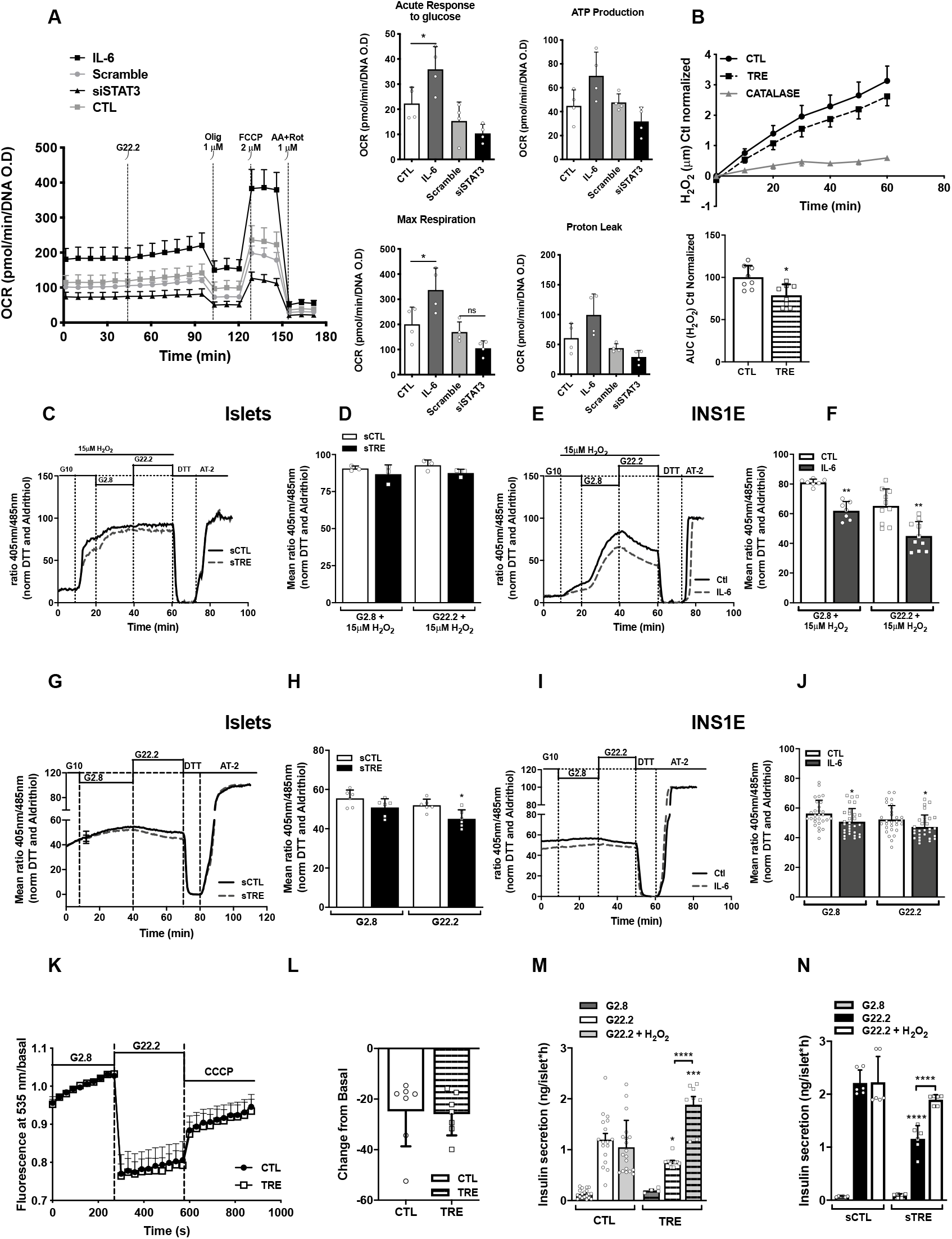
Exercise modulates beta cell function through IL-6 and pS-STAT3: Oxygen consumption rate (OCR) profiles of respective controls, IL-6-treated and STAT3 knocked down INS-1E cells in basal conditions (2.8 mM glucose) and after sequential treatment with 22.2mM of glucose (G22.2), oligomycin (1 μM), FCCP (2 μM) and rotenone plus antimycin A (1 μM each). In the Right panels: response to high glucose (after injection of 22 mM glucose), ATP production (calculated by subtracting the minimum measurement after oligomycin injection to the last measurement after glucose injection), maximal respiration (calculated by subtracting non-mitochondrial respiration to the maximum measurement after FCCP injection) and proton leak (calculated by subtracting non-mitochondrial respiration to the respiration after oligomycin injection) are shown (A). H2O2 production in islets from control (CTL) and trained (TRE) mice measured by amplex UltraRed probe (B). Dynamic fluorescent measurements in islets and INS-1E cells following transduction with cyto-roGFP1 or mito-roGFP1 (C-J). Islets treated during 24h with serum from control (sCTL) and trained (sTRE) mice expressing cyto–roGFP1 (C, D). INS-1E cells expressing cyto–roGFP1 treated during 48h with IL-6 (80ng/mL) (E, F). Islets treated per 24h with serum from control (sCTL) and trained (sTRE) mice expressing mito–roGFP1 (G, H). INS-1E cells treated per 48h with IL-6 (80ng/mL) expressing mito–roGFP1 (I, J). Islets and INS-1E cells were perfused with increasing concentrations of glucose followed by application of 10mM DTT and 100μM AT2 in the presence of glucose 10mM for normalization of the traces. ATP flow in islets from control (CTL) and trained (TRE) mice in response to glucose (K, L). GSIS in islets from control (CTL) and trained (TRE) mice in the presence of H2O2 (15μM) (M). GSIS in islets from untrained mice treated with serum from control (sCTL) and trained (sTRE) mice per 24h in the presence of H2O2 (5μM) (N). G2.8 represents 2.8mM and G22.2 represents 22.2mM of glucose. Values represent the mean ± SD of four to seven experiments. *P<0.05/**P<0.01/***P<0.001/****P<0.0001 vs. Control or as indicated.

### Obese trained mice displayed the same features of lean trained mice

The next step was to investigate whether prolonged training reduces insulin hypersecretion in a model of insulin resistance and obesity through the IL-6R signaling pathway. We subjected black 6 mice to an HFD leading to increased body fat, glucose intolerance, insulin resistance and insulin hypersecretion. Exercise training along with HFD, in addition to increasing fitness and performance levels (Table 2; P < 0.05), mitigated or prevented HFD effects by reducing body weight, fat content, glycemia, insulinemia, glucose intolerance, insulin resistance and GSIS (Figure 4 A-G; P < 0.05). Then, islets from lean, untrained mice were cultured with serum from obese trained (sHFDTRE) and obese untrained mice (sHFD), with or without TCZ. TCZ treatment abolished the training effect on GSIS (Figure 4H; P = 0.019). Moreover, INS-1E cells cultured with sHFDTRE showed higher amounts of pS-STAT3, and this increase was inhibited by exposure to TCZ (Figure 4I, P = 0.0615). As observed in islets from lean mice, exposure to exogenous H_2_O_2_ prevented the effects of training on GSIS down-regulation in islets from obese trained mice (Figure 4J; P = 0.0111).

**Table 2:** Functional adaptations to the training protocol in obese C57Bl/6 mice.

|  | **HFD** | **HFD TRE** |
| --- | --- | --- |
| **Body Weight (g)** | 30.40±0.95 | 25.60±0.72** |
| **VO<sub>2</sub>max. (ml.kg.min)** | 48.58±1.91 | 55.84±0.80* |
| **Distance (cm)** | 12077±1640 | 29804±1990* |
| **Maximum Speed (cm/s)** | 27.67±1.5 | 40.65±1.7* |

**Figure 4.**
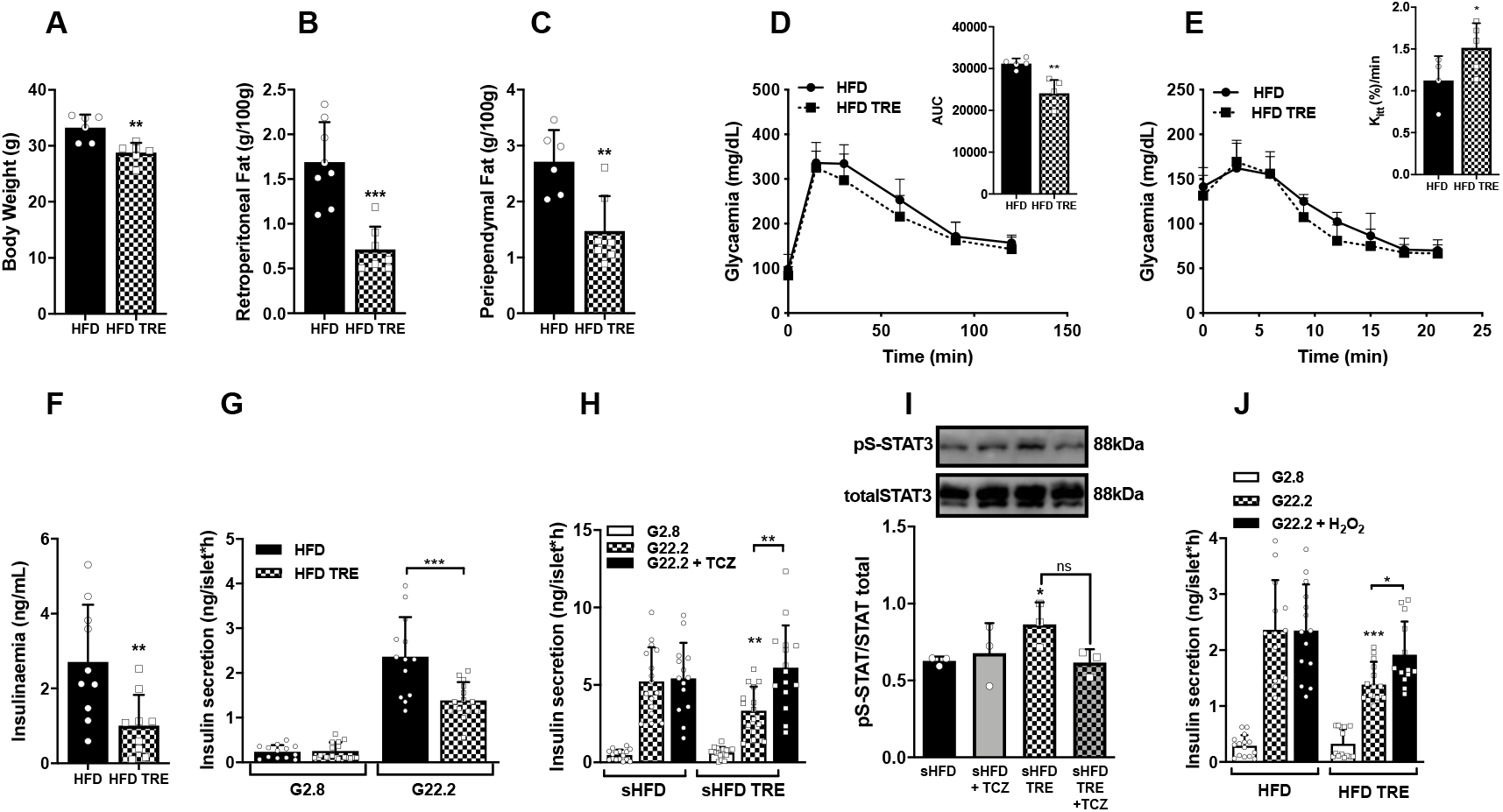
Exercise modulates beta cell function through pS-STAT3 in obese mice: Black 6 mice were placed under a high fat diet and remained untrained (HFD) or followed the endurance training protocol (HFD TRE). Body and fat pads weights (A-C). Glucose tolerance test (D). Insulin tolerance test (E) and insulinemia (F). Glucose-stimulated insulin secretion (GSIS), in response to 2.8mM (G2.8) or 22mM (G22.2) of glucose (G). GSIS in islets treated during 24h with mice serum and TCZ (100ng/mL) (H). Protein content pS-STAT3 in islets treated with serum plus TCZ (100ng/mL) (I). GSIS in the presence of H2O2 (15μM) (J). Values represent the mean ± SD of four to five experiments. *P<0.05/**P<0.01/***P<0.001 vs. Control or as indicated.

## DISCUSSION

We have previously reported a significant increase in serum IL-6 levels 1h after the end of the exercise, with IL-6 release peaking after 2–6 h, and returning to basal levels after 24h (Paula *et al*., 2015). The same exercise protocol was used in this study. The present findings suggest that the IL-6R/pS-STAT3 signaling axis modulates mitochondrial ETC activity and redox state in beta cells, contributing to exercise-induced GSIS down-regulation.

*In vivo* and *in vitro* experiments showed exercise leads to a decrease in GSIS, and this effect is prevented by IL-6R blockade. Previous studies investigating the impact of IL-6 peptide on beta cell function reported controversial results, showing increased as well as decreased GSIS (Ellingsgaard *et al*., 2011; Kristiansen and Mandrup-Poulsen, 2005). IL-6 is released in response to different metabolic conditions, and the source, milieu, triggering signal, and amount of IL-6 released result in diverse signaling effects (Trayhurn, Drevon and Eckel, 2011). These differences may account for the opposite effects of IL-6 observed on beta cell function. Ellingsgaard and colleagues showed that after an acute bout of exercise until exhaustion, IL-6 stimulates increased GSIS through an independent mechanism, involving GLP1 direct signaling to beta cells (Ellingsgaard *et al*., 2011). In agreement, unpublished experiments from our group reported higher levels of GSIS after acute bouts of endurance and strength exercises. Interestingly, each exercise stimulus leads to different effects on GSIS, which remain to be further investigated.

Here, we investigated the effects of a chronic exercise training regimen, which lasted 8 weeks, on reducing GSIS. In this sense, the reestablishment of GSIS after IL-6R blockade and STAT3 knock-down provide strong evidence for chronic exercise signaling to beta cells through the IL-6R/pS-STAT3 axis. On the other hand, we show that IL-6 concentration in the serum of trained mice, at the point of blood sample collection (48h after last exercise session), is lower compared to CTL. These results suggest the possibility of other IL-6R ligands rather than or in addition to IL-6, as likely triggering molecules of this signaling under physiological conditions. Moreover, the high IL-6 concentration needed to mimic experiments using sera from trained mice outcomes reinforces this hypothesis. Further studies are required to elucidate the complex IL-6R and its signaling pathway activation on exercise-induced beta cell function.

In the next step, we evaluated whether exercise elicits STAT3 non-canonical signaling in beta cells. Our results showed higher pS-STAT3 content in the mitochondrial fraction of INS-1E cells after exposure to IL-6. Additionally, the fact that TCZ prevents mitochondrial pS-STAT3 increase reinforced the possibility of the activation of non-canonical IL-6R signaling triggered by exercise. Studies investigating pS-STAT3 signaling showed it may bind to the complexes I, II or V, increasing ETC activity and reducing mitochondrial ROS production (Gough *et al*., 2009; Heusch *et al*., 2011; Szczepanek *et al*., 2011; Wegrzyn *et al*., 2009). In agreement with the aforementioned studies and corroborating the pS-STAT3 mitochondrial location also in the beta cells, our results showed that pS-STAT3 binds to ATP synthase (complex V). Accordingly, functional experiments showing higher oxygen consumption support the molecular data and provide primary evidence for increased ETC activity in response to IL-6R signaling-induced mitochondria STAT3 translocation. This outcome was reinforced when cells lacking STAT3 displayed lower rates of oxygen consumption. The lower amount of mitochondrial ROS observed in islets and INS-1E cells cultured with exercise-conditioned serum or IL-6 provide additional evidence for the enhancement of ETC activity. The observed increase on INS-1E maximal oxygen consumption suggests pS-STAT3 might induce a mild uncoupling, sufficient to reduce ROS production, establishing a new redox set point, leading to GSIS down-regulation. In line with our findings, it was previously shown in CD4^+^ T cells that IL-6 signaling plays a pivotal role in the induction of pS-STAT3 translocation to the mitochondria. The same study also showed higher ETC activity and consequent decrease of mitochondrial ROS production in response to IL-6 exposure (Yang *et al*., 2015). To investigate the likely mechanistic link between increased ETC activity-induced lower ROS production and GSIS down-regulation, we exposed islets to exogenous H_2_O_2_, leading to the loss of exercise effect on GSIS down-regulation.

To evaluate whether these outcomes would be reproduced in an insulin hypersecretion condition, we studied a model of diet-induced obesity. Exercise training prevented insulin hypersecretion in mice fed with a HFD. Moreover, the major mechanistic findings described above were reproduced in this insulin hypersecretion model.

In summary, although our results provide evidence that the IL-6R/STAT3 axis exerts a role on GSIS down-regulation, we cannot disregard the participation of other circulating factors, instead or in addition to IL-6, on the activation of this signaling pathway. Our findings reinforce the role of this pathway on the signal transduction of exercise benefits to the beta cells (Figure 5), highlighting this axis as a potential therapeutic target for the prevention and/or treatment of metabolic diseases associated with hyperinsulinism and beta cell failure.

**Figure 5.**
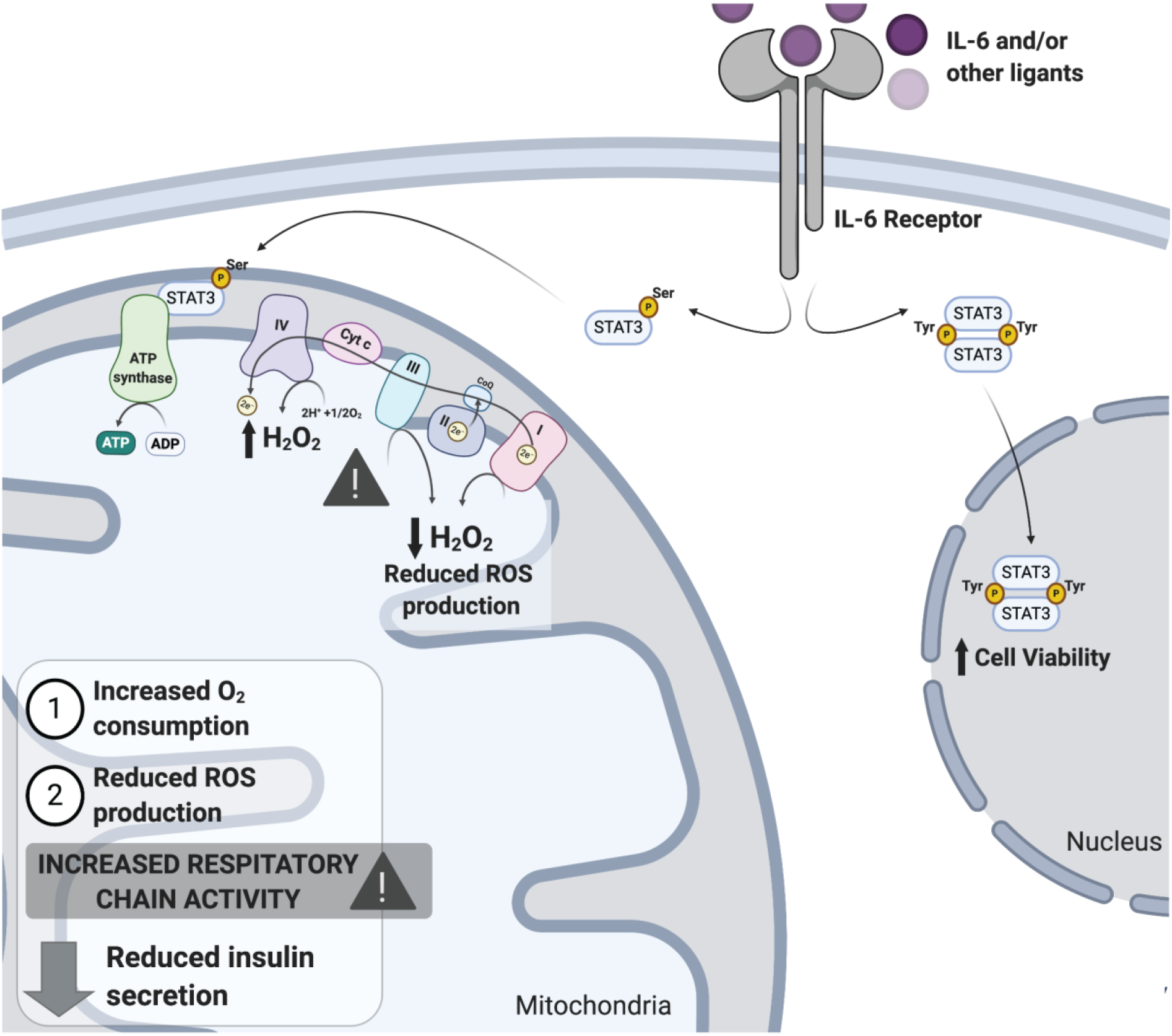
Proposed schematic representation of exercise-induced GSIS modulation through IL-6/STAT3 pathway signaling: IL-6 is released in response to exercise training, binds to its receptor in the beta cells and signals STAT3 phosphorylation at tyrosine 705 and serine 727 residues. STAT3 phosphorylation at the serine 727 residue enables its translocation to the mitochondria and its binding to the ATP synthase (complex V). This interaction promotes an increase in the ETC activity and reduces ROS content. Lower ROS levels lead to insulin secretion down-regulation.

## ADDITIONAL INFORMATION

## Acknowledgments

The authors acknowledge the financial support of FAPESP grants: 2014/01717-9 and 2015/12611-0 to A.C.B., E.M.C. and C.C.Z., 2013/00750-0 to N.C.L. and “Allocation de recherche 2016” from the Société Francophone du diabète to J.J., J.J. is Research Director from the Fonds de la Recherche Scientifique-FNRS, Belgium.

## Author contributions

N.C.L., C.C.Z., and E.M.C. designed the study; N.C.L., F.P., C.L., P.C.B., J.J.M., and J.C.S. performed the experiments and analyzed the data; N.C.L, F.M., C.L., C.C.Z, and J.J. interpreted the data; E.M.C., D. L. E., J.J., and A.C.B. provided materials for this study; N.C.L. and C.C.Z. wrote the manuscript; C.C.Z., E.M.C., J.J. and D.L.E. supervised the study. All authors reviewed the manuscript.

## Data availability statement

The data that support the findings of this study are available from the corresponding author upon reasonable request.

