## Supplementary Information for "Non-canonical mitochondrial STAT3 signaling mediates exercise-induced insulin secretion down-regulation"

[1]Obesity and Comorbidities Research Center (OCRC), Department of Structural and Functional Biology, Institute of Biology, University of Campinas (UNICAMP), Campinas, Brazil.
[2]Université Catholique de Louvain, Institute of Experimental and Clinical Research, Pole of Endocrinology, Diabetes and Nutrition, Brussels, Belgium.
[3]ULB Center for Diabetes Research, Université Libre de Bruxelles (ULB), Brussels, Belgium.
[#]These authors contributed equally to the study
*Senior co-authorship

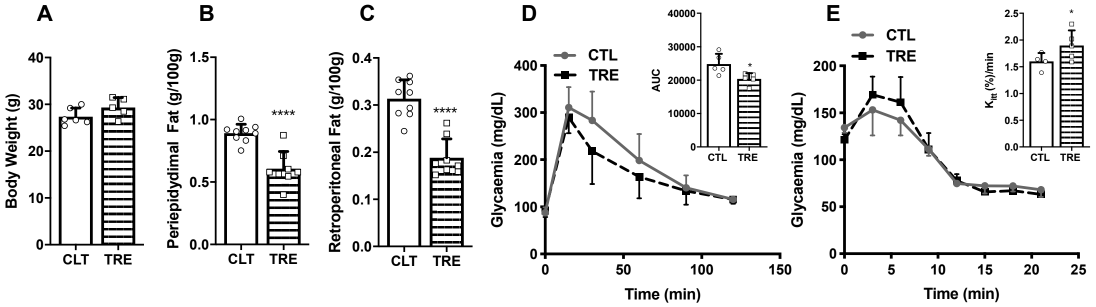


*<u>Supplementary figure 1</u>. Trained mice present improvement of glucose homeostasis: Body weight (A). Fat Pads (B, C). Glucose tolerance test and corresponding area under the curve (AUC) (D). Insulin tolerance test and glucose decay constant rate (Kitt) (E). Values represent the mean ± SD of three to four independent experiments. *P<0.05/**P<0.01/***P<0.001/**** P<0.0001 vs. Control or as indicated.*

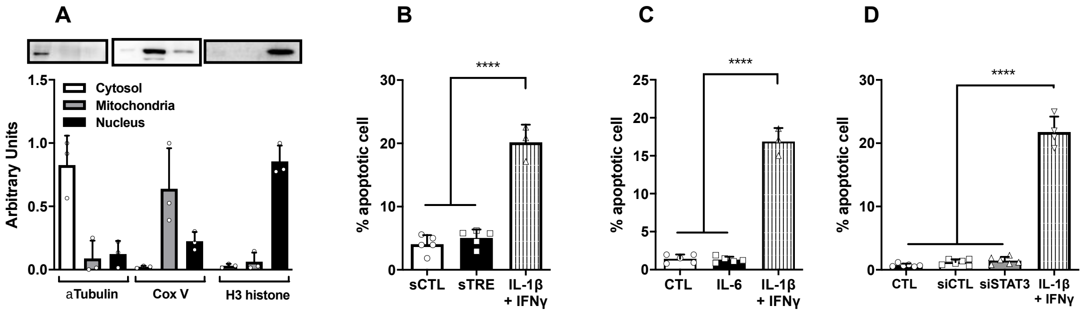


*<u>Supplementary figure 2</u>. Validation of fraction purity measuring specific cellular compartments proteins (A). Fluorescence quantification of INS-1E cells stained for HO-PI treated with serum from control (sCTL) and trained (sTRE) mice (B), IL-6 (C) or siSTAT3 (D). Values represent the mean ± SD of three to eleven experiments. *P<0.05/**P<0.01/***P<0.001/****P<0.0001 vs. Control or as indicated.*

### Uncropped blots used in this study

**Figure 1L**

Membrane 01

totalSTAT3

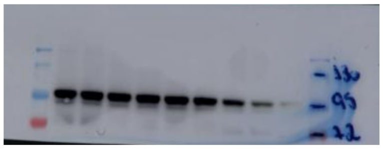

Membrane 01

αTubulin

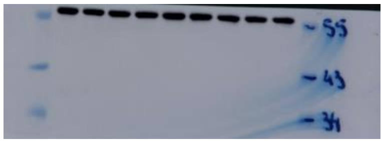

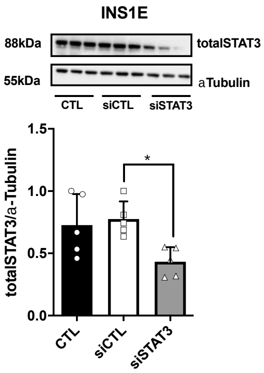


**Figure 2A**

Membrane 02

pS-Stat3

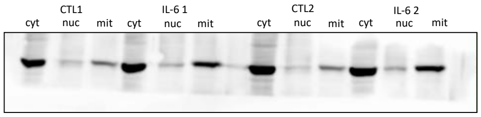


Membrane 02

totalSTAT3

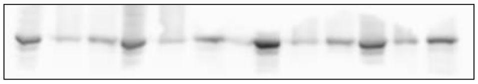

Membrane 03

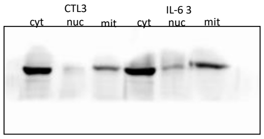


Membrane 03

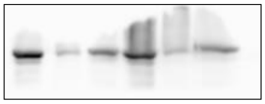

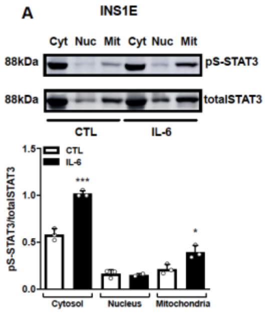

**Figure 2B**

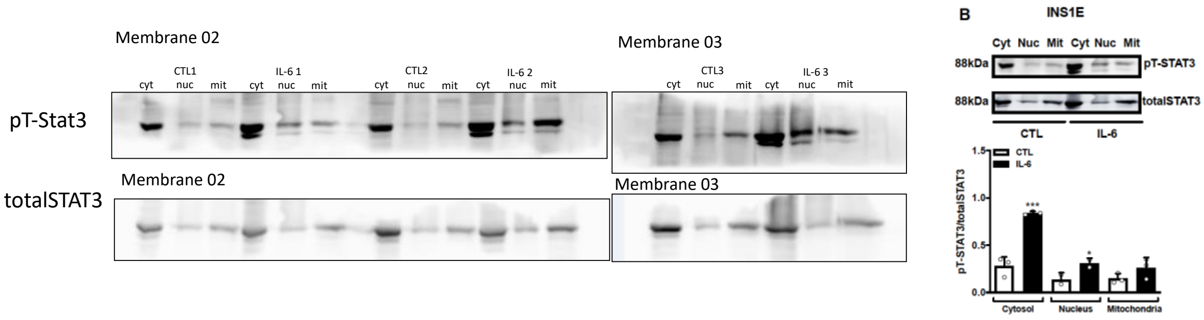


**Figure 2C**

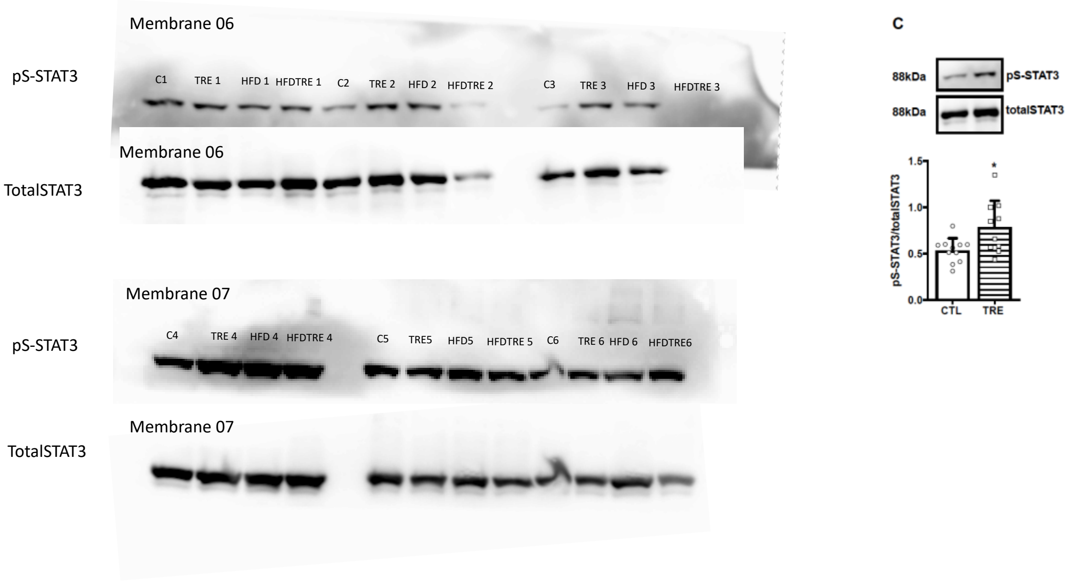

**Figure 2D**

pS-STAT3

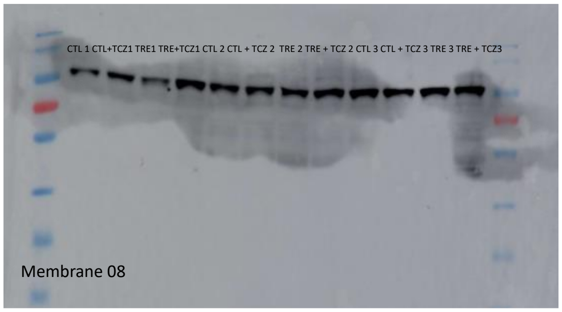


TotalSTAT3

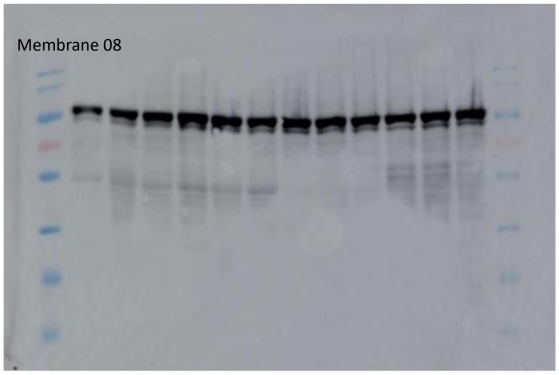


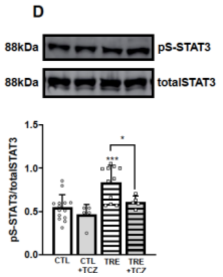


**Figure 2E**

pS-STAT3

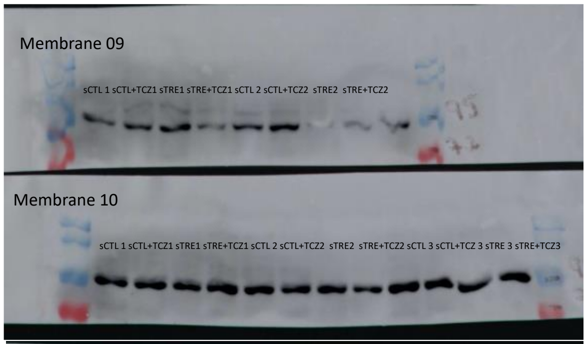


TotalSTAT3

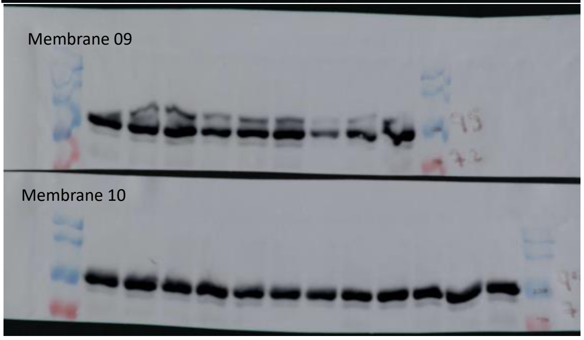


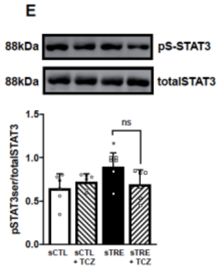

### Figure 4I

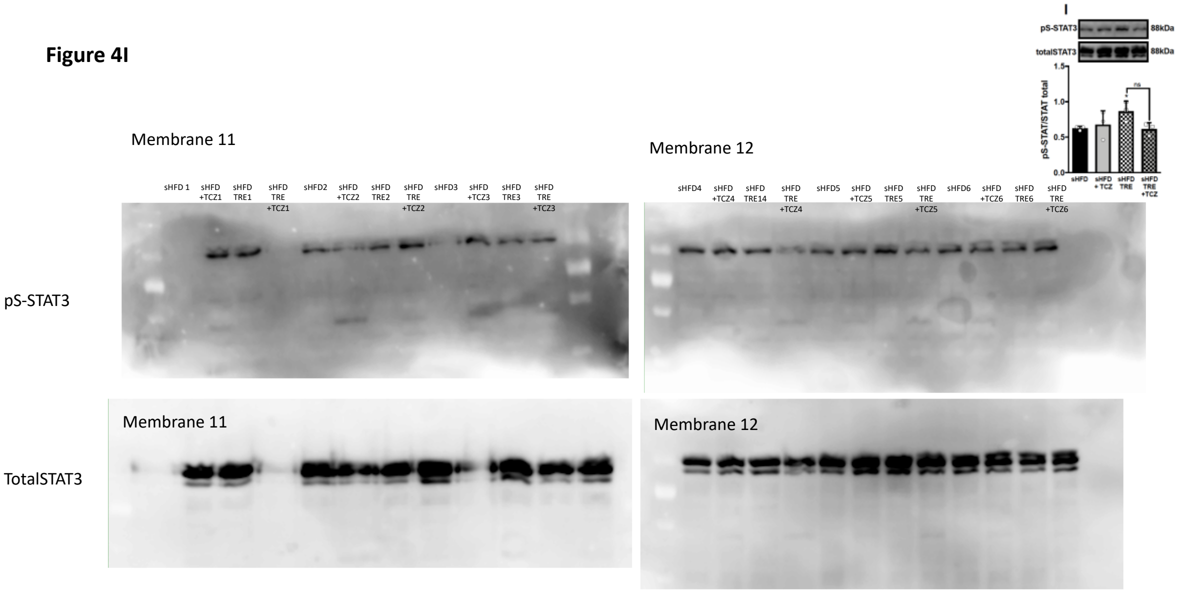
I
pS-STAT3
88kDa
totalSTAT3
88kDa
pS-STAT/STAT total
ns
sHFD
sHFD + TCZ
sHFD TRE
sHFD TRE +TCZ
Membrane 11
Membrane 12
sHFD 1
sHFD +TCZ1
sHFD TRE1
sHFD TRE +TCZ1
sHFD2
sHFD +TCZ2
sHFD TRE2
sHFD TRE +TCZ2
sHFD3
sHFD +TCZ3
sHFD TRE3
sHFD TRE +TCZ3
sHFD4
sHFD +TCZ4
sHFD TRE14
sHFD TRE +TCZ4
sHFD5
sHFD +TCZ5
sHFD TRE5
sHFD TRE +TCZ5
sHFD6
sHFD +TCZ6
sHFD TRE6
sHFD TRE +TCZ6
pS-STAT3
Membrane 11
Membrane 12
TotalSTAT3
